## Supplemental Figures for Manuscript for "Triple Negative Breast Cancer Cells Acquire Lymphocyte Proteins and Genomic DNA During Trogocytosis with T Cells"

Science Advances
AAAS

### Supplementary Materials for

#### Triple Negative Breast Cancer Cells Acquire Lymphocyte Proteins and Genomic DNA During Trogocytosis with T Cells

Anutr Sivakoses *et al.*

**This PDF file includes:**

Figs S1 to S4

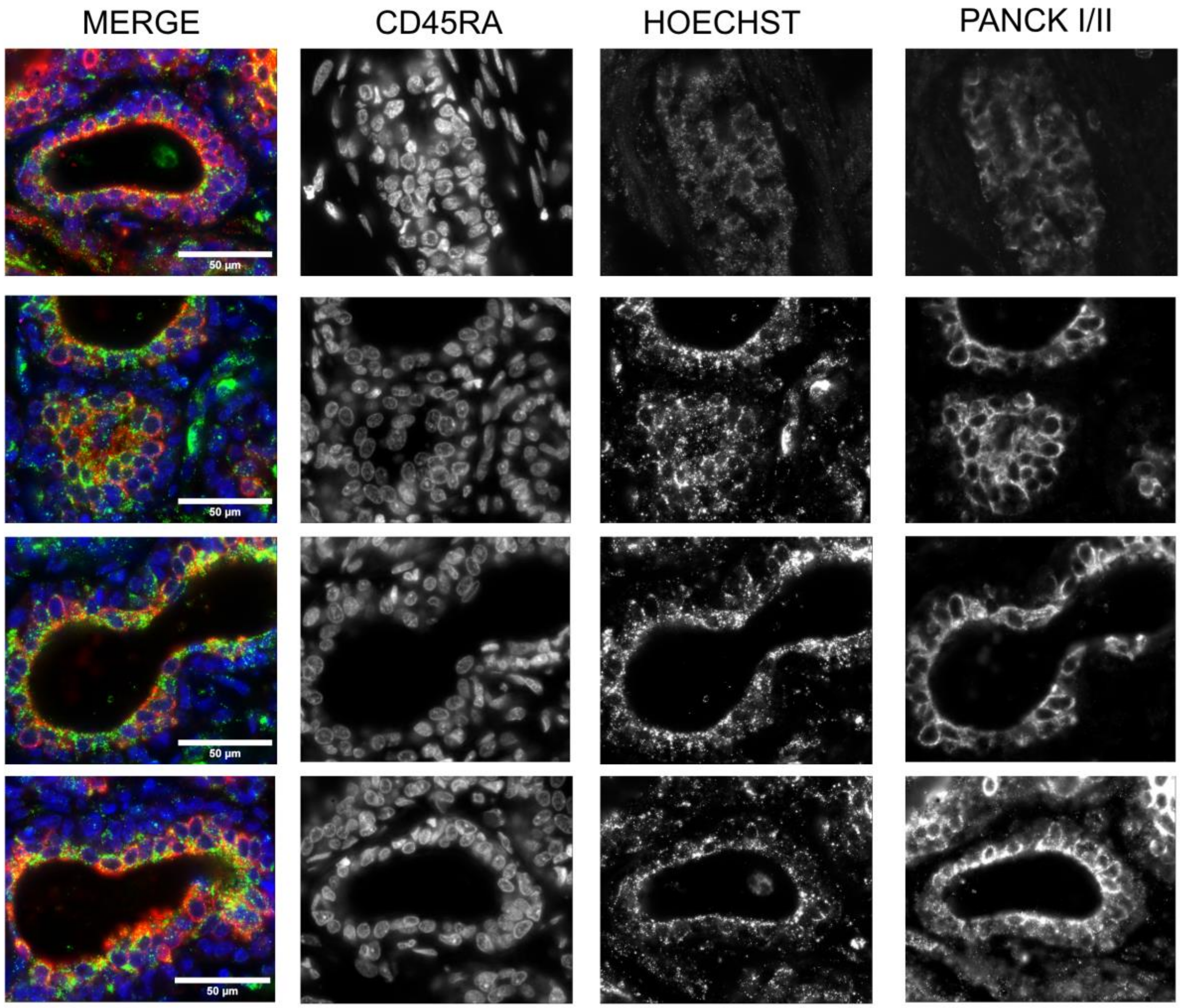


**Figure S1.** Additional Images of TNBC cells expressing CD45RA. Merge images are shown on the left panel with single channel images for CD45RA, Hoechst, and tumor marker Pan-Cytokeratin Type I/II shown in their respective columns.

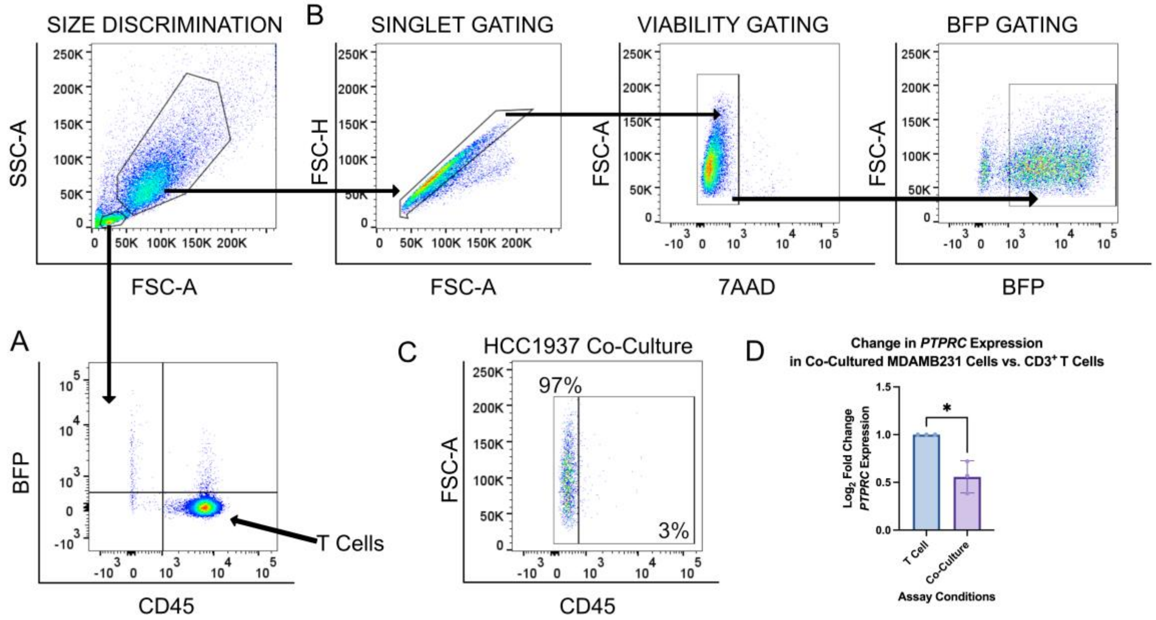


**Figure S2.** Flow Cytometry analysis of TNBC cell lines. (A) Analysis of T cells to visualize lack of BFP-LA acquisition post co-culture. (B) Gating strategy to identify viable, BFP-LA$^+$ tumor singlets. (C) Co-culture analysis of HCC1937 TNBC cells (D) Change in expression of *PTPRC* between co-cultured MDAMB231 and primary T cells. An unpaired T test was used to perform statistical analysis on this data *p = 0.0104.

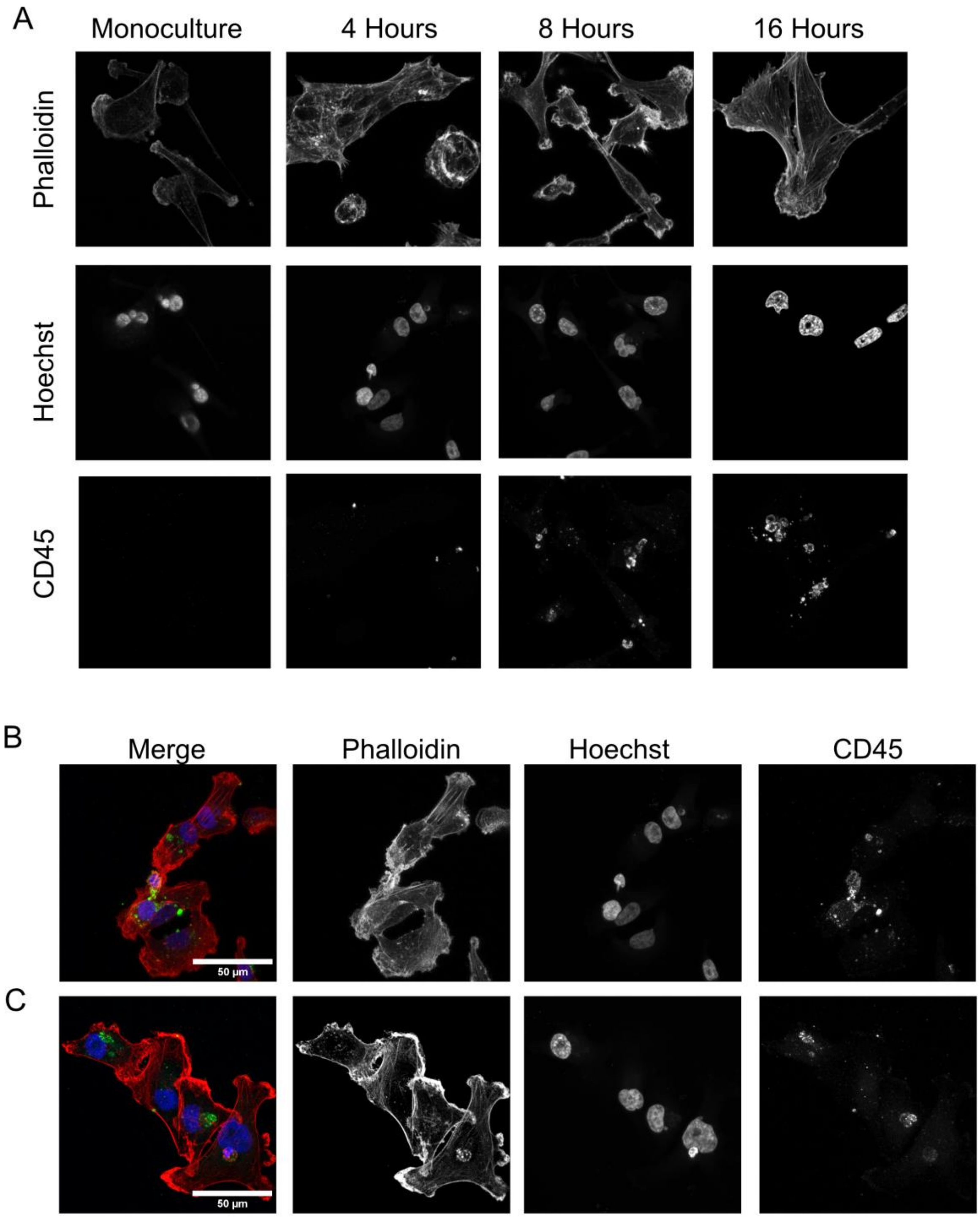
A
Monoculture
4 Hours
8 Hours
16 Hours
Phalloidin
Hoechst
CD45
B
Merge
Phalloidin
Hoechst
CD45
50 µm
C
50 µm

**Figure S3.** (A) Single channel images of co-cultured MDAMB231 shown in Fig. 3 and additional merge and single channel images of trogosomes found present in (B) MDAMB231 and (C) MDAMB436 cells.

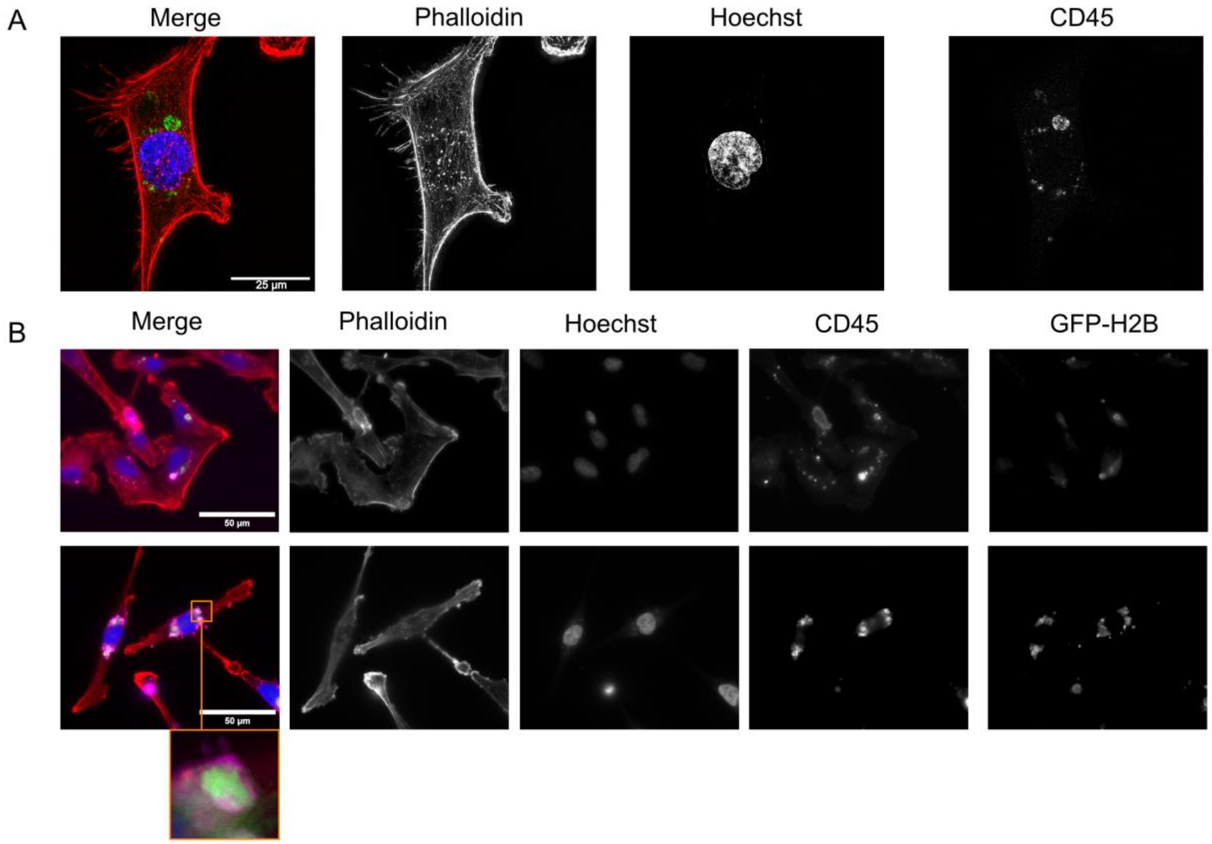


**Figure S4.** Additional images of TNBC cells acquiring genomic DNA from T cells. (A) SIM image of MDAMB231 cells containing a trogosome without positive Hoechst labeling. (B) Images of MDAMB231 cells containing GFP-H2B with merge images on the leftmost panel and single channel images in their respective columns.
